# Proteomic Differences in the Hippocampus and Cortex of Epilepsy Brain Tissue

**DOI:** 10.1101/2020.07.21.209163

**Authors:** Geoffrey Pires, Dominique Leitner, Eleanor Drummond, Evgeny Kanshin, Shruti Nayak, Manor Askenazi, Arline Faustin, Daniel Friedman, Ludovic Debure, Beatrix Ueberheide, Thomas Wisniewski, Orrin Devinsky

## Abstract

Epilepsy is a common neurological disorder affecting over 70 million people worldwide, with a high rate of pharmaco-resistance, diverse comorbidities including progressive cognitive and behavioral disorders, and increased mortality from direct (e.g., Sudden Unexpected Death in Epilepsy [SUDEP], accidents, drowning) or indirect effects of seizures and therapies. Extensive research with animal models and human studies provides limited insights into the mechanisms underlying seizures and epileptogenesis, and these have not translated into significant reductions in pharmaco-resistance, morbidities or mortality. To help define changes in molecular signaling networks associated with epilepsy, we examined the proteome of brain samples from epilepsy and control cases. Label-free quantitative mass spectrometry (MS) was performed on the hippocampal CA1-3 region, frontal cortex, and dentate gyrus microdissected from epilepsy and control cases (n=14/group). Epilepsy cases had significant differences in the expression of 777 proteins in the hippocampal CA1-3 region, 296 proteins in the frontal cortex, and 49 proteins in the dentate gyrus in comparison to control cases. Network analysis showed that proteins involved in protein synthesis, mitochondrial function, G-protein signaling, and synaptic plasticity were particularly altered in epilepsy. While protein differences were most pronounced in the hippocampus, similar changes were observed in other brain regions indicating broad proteomic abnormalities in epilepsy. Among the most significantly altered proteins, G-protein Subunit Beta 1 (GNB1) was one of the most significantly decreased proteins in epilepsy in all regions studied, highlighting the importance of G-protein subunit signaling and G-protein–coupled receptors (GPCRs) in epilepsy. Our results provide insights into the molecular mechanisms underlying epilepsy, which may allow for novel targeted therapeutic strategies.

## Introduction

Epilepsy is a spectrum disorder with diverse etiologies and outcomes affecting over 70 million people worldwide [1–4]. Etiologies include ion channel and many other genetic mutations, infection, structural lesions, metabolic abnormalities, and autoimmune disorders, although the cause remains unknown in many cases [5–9]. The mechanisms underlying ictogenesis and epileptogenesis alter the balance of excitation and inhibition [10]. Anti-seizure drugs (ASDs) target multiple mechanisms, including ion channels (e.g., sodium, calcium, potassium), neurotransmitter receptors and metabolism (e.g., GABA-A receptors, GABA transporter 1, GABA transaminase), synaptic vesicles (e.g., SV2A) and regulators of cell growth (e.g., mTOR inhibition). However, since a third of patients have an incomplete response to ASDs, dietary, neuromodulatory and resective or disconnective surgical procedures, a broader understanding of the cellular and molecular drivers of ictogenesis and epileptogenesis is essential to identify new therapeutic targets. [4, 11, 12].

Human and animal epilepsy studies reveal that seizures can arise from neocortical or limbic lesions, including focal cortical dysplasia (FCD), hippocampal sclerosis, tumors, inflammation and vascular lesions [13–15]. The hippocampus is often involved in focal epilepsy as a primary (e.g., mesial temporal lobe epilepsy [MTLE]) or secondary seizure focus (e.g. areas robustly connected with entorhinal cortex and hippocampus). The hippocampal Cornu Ammonis (CA) CA1-4 region and dentate gyrus are involved in seizure generation and in hippocampal sclerosis [16].

To date, our knowledge of brain protein changes in epilepsy comes from experimental MTLE models and human brain tissue. The most robust changes in epilepsy brain tissue involve proteins related to neurotransmitters, cell signaling, glucose metabolism and protein synthesis [17, 18]. Genomic, epigenetic and transcriptomic studies have revealed insights into mechanisms underlying ictogenesis and epileptogenesis by identifying novel genetic loci and profound changes in gene expression in humans and animal models [19–24]. However, the poor correlation between RNA expression and protein levels limits the value of isolated transcriptomic or genomic studies [25]. Unbiased, mass spectrometry-based proteomic studies of human epilepsy tissue can complement these studies, particularly since proteins are direct drug targets. Proteomics provides a global view of protein changes at a functional or network level. The majority of previous epilepsy proteomics studies have examined animal model tissue, rather than human brain tissue [26, 27]. Two previous studies have performed proteomics on human epilepsy brain tissue, however these were limited by small numbers, analysis of single regions and low sensitivity proteomics approaches. These studies identified 77 proteins differentially expressed in MTLE hippocampi compared to controls [28] and 18 differentially expressed proteins between high and low spiking regions in six epilepsy patients [29]. We hypothesized that we would identify more extensive protein differences in epilepsy by using a more sensitive proteomic approach, and that we would generate novel insight into the pathophysiology of epilepsy by examining protein changes in multiple brain regions that are differentially affected in epilepsy.

Therefore, to advance our knowledge of proteomics in human epilepsy, we used label-free quantitative mass spectrometry (MS) in epilepsy and control human brains across three regions: CA1-3 of the hippocampus, the dentate gyrus and the frontal cortex.

## Methods

### Ethics Statement

All studies were approved by the Institutional Review Board (IRB) at New York University (NYU). In all cases, written informed consent was obtained from the legal guardian or patient, and material used had ethical approval. All patient data and samples were deidentified and stored according to NIH guidelines.

### Human Brain Tissue

All human tissue samples were collected at autopsy. Epilepsy and control cohorts were studied. The epilepsy cases (n=14) were from the North American SUDEP Registry (NASR) and cause of death was not SUDEP. The control cases (n=14) were from the Center for Biospecimen Research and Development (CBRD) at NYU. Histological evaluation of all cases was performed by neuropathologists and findings were classified using the international consensus criteria [2, 13, 30]. Several cases did not have comprehensive clinical information, which came from a population with health care disparities that were not under regular medical care and therefore had limited medical records. Regarding epilepsy syndrome, there was a broad representation of various epilepsies in the cases with known clinical information, the majority of which excluded known frontal lobe involvement. All cases were matched for sex and age (46 ± 14 years) to the best of our ability. Individual patient information (sex, age, cause of death (COD), post mortem interval (PMI) and seizure classification if applicable) is included in Table 1.

**Table 1.** Control and Epilepsy Case Information.

| Group | Age | Sex | COD | PMI | Significant Neuropathology | Seizure Classification | Brain Region |
| --- | --- | --- | --- | --- | --- | --- | --- |
| **<u>Controls</u>** |  |  |  |  |  |  |  |
| 1 | 55 | M | Cardiac arrest | 64 |  |  | HP, DG, FC |
| 2 | 57 | M | Cardiac arrest | 142 |  |  | HP, DG, FC |
| 3 | 56 | F | Cardiac arrest | 46 |  |  | HP, DG, FC |
| 4 | 49 | F | Cardiac arrest | 144 |  |  | HP, DG, FC |
| 5 | 59 | F | Cardiac arrest | 57 |  |  | HP, DG, FC |
| 6 | 59 | M | Possible cardiac arrest | 116 |  |  | HP, DG, FC |
| 7 | 57 | M | Cardiac arrest | 91 |  |  | HP, DG, FC |
| 8 | 55 | M | Pulmonary fibrosis | 57 | Subacute, right temporal white matter microinfarct |  | HP, DG, FC |
| 9 | 59 | M | Cardiac arrest | 60 |  |  | HP, DG, FC |
| 10 | 38 | M | Anoxic encephalopathy secondary to cardiac arrest | 44 | Acute and chronic hypoxic-ischemic changes, diffuse |  | HP, DG, FC |
| 11 | 59 | M | Supraglottic squamous cell carcinoma | <120 | Ischemic infarct, left frontal MCA/ACA |  | HP, DG, FC |
| 12 | 49 | M | Cardiac arrest | 48 | Meningeal fibrosis |  | HP, DG, FC |
| 13 | 50 | M | Possible cardiac arrest (contributing - acute gastroenteritis) | 50 | Hypoxic-ischemic changes, diffuse |  | HP, DG, FC |
| 14 | 54 | M | Cardiac arrest | 66 | Hypoxic-ischemic injury, acute, multifocal |  | HP, DG, FC |
| **<u>Epilepsy</u>** |  |  |  |  |  |  |  |
| 1 | 36 | M | Overdose/intoxication | 20 |  | Unclassified | HP, DG, FC |
| 2 | 54 | M | Accident/trauma | <24 | Mild gliosis, contusion, disorganization | ND | HP, DG |
| 3 | 64 | F | Overdose | 18 |  | Generalized, Unclassified | HP, DG |
| 4 | 50 | M | Choking on foreign object | 15 |  | Focal | FC |
| 5 | 9 | F | Drowning | 30 | FCD | ND | HP, DG |
| 6 | 45 | M | Suicide | 27 | Dysgenesis | Focal | HP, DG |
| 7 | 36 | M | Drowning | 48 | Sclerosis | Focal | HP, DG, FC |
| 8 | 45 | M | Suicide | <48 |  | Unclassified | HP, DG, FC |
| 9 | 24 | F | Drowning | <48 | Dysgenesis | ND | HP, DG, FC |
| 10 | 28 | M | Accident/trauma | <48 | Dysgenesis | Unclassified | HP, DG, FC |
| 11 | 23 | M | Drowning | <48 | FCD | Unclassified | HP, DG, FC |
| 12 | 34 | F | Pulmonary embolism | 13 | FCD | Focal | HP, DG, FC |
| 13 | 32 | M | Ethanol intoxication and clobazam overdose | 19 | FCD, Wernicke's encephalopathy | ND | HP, DG, FC |
| 14 | 49 | M | Aspiration | 43 | Dysgenesis, sclerosis,<br>gliosis, hemisphere<br>atrophy | Unclassified | HP, DG, FC |

PMI = post-mortem interval, COD = cause of death, MCA/ACA = middle cerebral artery/anterior cerebral artery, FCD = focal cortical dysplasia, ND = not determined, HP = hippocampus, DG = dentate gyrus, FC = frontal cortex

### Laser-Capture Microdissection (LCM)

We sampled brain tissue from formalin-fixed paraffin embedded (FFPE) blocks of hippocampus or superior frontal gyrus collected and processed for autopsy neuropathology. LCM was performed using our published protocol [31–33]. Briefly, 8μm-thick FFPE sections containing hippocampus or frontal superior gyrus were collected onto LCM PET Frame Slides (Leica). Sections were stained with cresyl violet to visualize neurons and regions of interest (ROIs). For each sample, 10mm^2^ of hippocampus (including CA1-3) and superior frontal gyrus (including layers I-IV) and 4mm^2^ of dentate gyrus (granule cell layer) were microdissected in separate tubes using a LMD6500 microscope at 5X magnification (Leica) (Figure 1). ROIs were collected into ultrapure Pierce Water, LC-MS Grade (Thermo Scientific). After collection, samples were centrifuged at 14,000g for 2min and stored at −80°C until peptide extraction.

**Figure 1.**
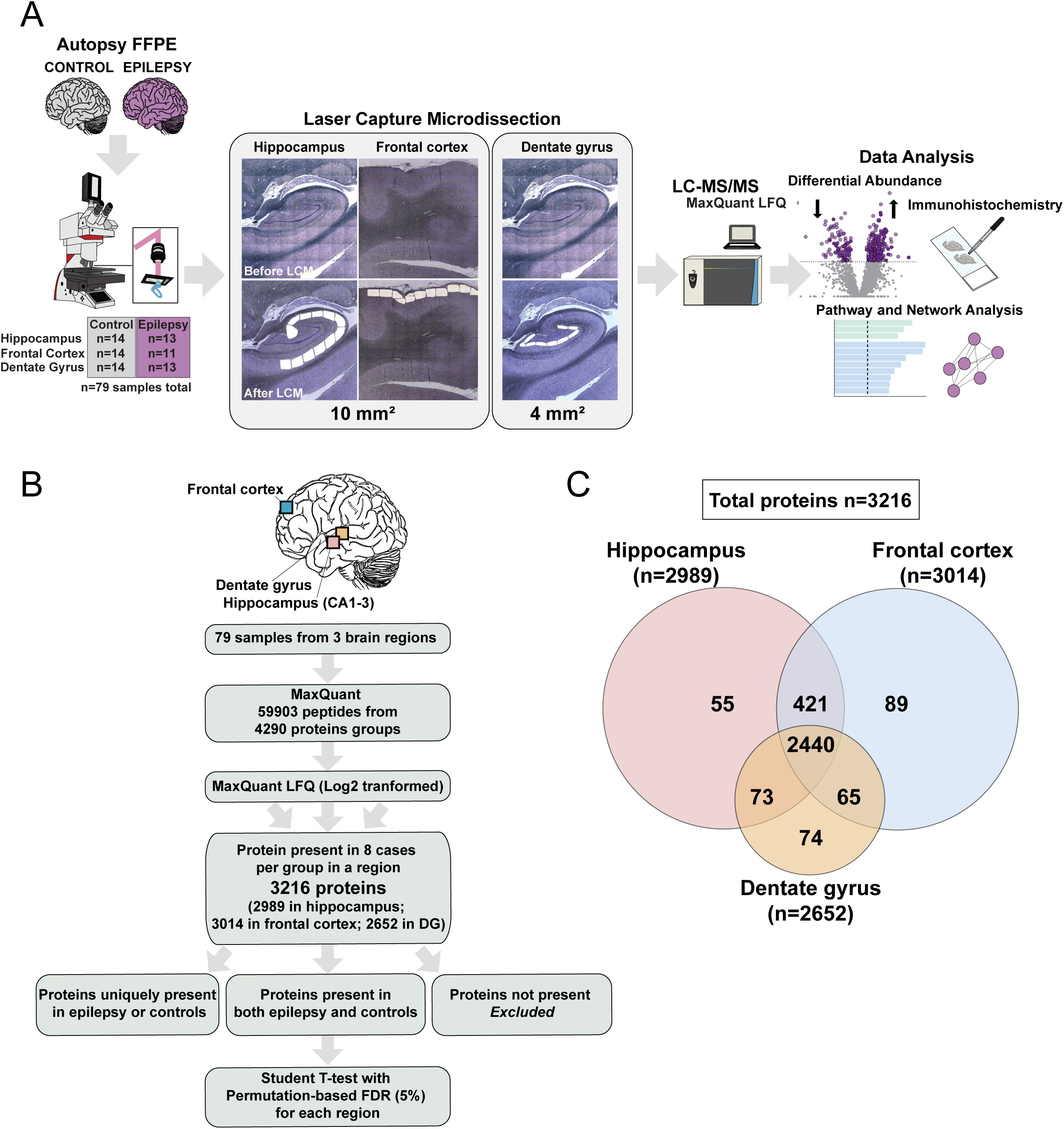
Quantitative proteomics analysis of epilepsy and non-epilepsy control cases. **A. Experimental workflow.** Specific protein regions were excised by Laser Capture Microdissection (LCM) from FFPE autopsy brain tissue from control and epilepsy cases (n=14 in each group) and proteins quantified by label-free quantitative mass spectrometry (MS). Statistical, pathway and network analyses were performed to identify proteins differing in these regions. Three regions were analyzed (Hippocampus CA1-3 (n=14 control, n=13 epilepsy), Frontal Cortex (n=14 control, n=11 epilepsy) and Dentate gyrus (n=14 control, n=13 epilepsy). Validation of protein differences was performed using immunohistochemistry (IHC). **B. LC-MS/MS computational analysis.** Proteins groups were filtered and quantified using MaxQuant. **C. Venn diagram of proteins identified in each region.** A total of 3216 protein groups were detected across all brain regions by MS, with 2440 overlapping protein groups between the three regions and 2861 overlapping protein groups between the hippocampus and the frontal cortex.

### Label-free Quantitative Mass Spectrometry (MS) Proteomics

The study included 6 experimental groups based on two states (epilepsy, control) and three brain regions. Samples were processed and analyzed using label-free quantitative MS in three batches: hippocampus (n=13 epilepsy, n=14 controls), frontal cortex (n=11 epilepsy and n=14 control) and dentate gyrus (n=13 epilepsy, n=14 control). The same epilepsy and control cases were used in the three batches. For each ROI analyzed, samples were prepared for label-free quantitative MS [31, 32]. Samples were thawed at room temperature (RT) and resuspended by adding 50 μl of 100mM Ammonium Bicarbonate and 20% acetonitrile solution to the cap and the tube spun down. This step was repeated three more times to ensure thorough cap washing. Samples were then incubated at 95°C for 1hr followed by a 65°C incubation for 2hrs. Reduction was performed using dithiothreitol at 57 °C for 1hr (2 μl of 0.2 M) followed by alkylation with iodoacetamide at RT in the dark for 45min (2 μl of 0.5 M). The samples were subsequently digested with 300ng of sequencing grade modified trypsin (Promega) overnight at RT with gentle shaking. After digestion samples were acidified with trifluoroacetic acid to a pH of 2 and peptides were desalted using POROS R2 C18 beads as previously described [31–33]. The samples were resuspended in 0.5% acetic acid and stored at −80°C until further analysis. Aliquots of the sample were loaded onto an Acclaim PepMap 100 (75 um × 2 cm) precolumn connected to a PepMap RSLC C18 (2 um, 100A × 50 cm) analytical column using the autosampler of an EASY-nLC 1200 (Thermo Scientific). The samples were gradient eluted directly into an Orbitrap Fusion Lumos mass spectrometer (Thermo Scientific) using a 165min gradient (Solvent A consisting of 2% acetonitrile in 0.5% acetic acid and Solvent B of 80% acetonitrile in 0.5% acetic acid). High resolution full mass spectra (MS) were acquired with a resolution of 240,000, an automatic gain control (AGC) target of 1e6, a maximum ion time of 50ms, and a scan range of 400-1500 m/z. After each full MS, MS/MS HCD spectra were acquired in the ion trap. All MS/MS spectra were collected using the following: ion trap scan rate Rapid, AGC target of 2e4, maximum ion time of 18ms, one microscan, 2 m/z isolation window, fixed first mass of 110 m/z, and NCE of 32. The mass spectrometry raw files are accessible under MassIVE ID: MSV000085517.

### Proteomics Computational Analysis

MS data were analyzed using MaxQuant [34, 35] software version 1.6.3.4 and searched against the SwissProt subset of the *H. sapiens* Uniprot database (http://www.uniprot.org/) containing 20,365 entries (February 2019 release) to which 248 common laboratory contaminants were added. The enzyme specificity was set to trypsin with a maximum number of missed cleavages set to 2. Peptide identification was performed with an initial precursor mass deviation of 7 ppm and a fragment mass deviation of 20 ppm with subsequent non-linear mass recalibration [36]. Oxidation of methionine residues was searched as variable modification; carbamidomethylation of cysteines was searched as a fixed modification. The false discovery rate (FDR) for peptide, protein, and site identification was set to 1% and was calculated using a decoy database approach. The minimum peptide length was set to 6. The option match between runs (1min time tolerance) was enabled to correlate identification and quantitation results across different runs. Protein quantification was performed with built-in MaxQuant LFQ algorithm [37] with the following settings: minimum ratio count of 2, “fastLFQ” option enabled, minimum/average number of neighbors 3/6. LFQ normalization was performed within sample groups that belong to the same brain region.

For subsequent statistical analyses we limited our dataset to proteins quantified in at least 8 samples in at least one of the conditions/brain region. All common contaminant entries were removed, resulting in a matrix with 3216 protein groups. Data analysis used Perseus v. 1.6.2.3. [38] (http://www.perseus-framework.org/) using R environment for statistical computing and graphics (http://www.r-project.org/). Protein interaction networks and enrichment analyses were done in STRING [39, 40] v. 11.0 (https://string-db.org/).

### Statistical Analyses and Interpretation of Results

For Principal Component Analysis (PCA) missing values were imputed from a normal distribution with width 0.3 and downshift 1.8 using Perseus. Hierarchical clustering was used to visualize sample metadata in the context of a clustering those proteins associated with disease (whether positively or negatively, with a 5% FDR), following a preliminary kmeans clustering using k=2 and the Hartigan and Wong algorithm using the *ComplexHeatmap* package in the R environment [41]. For the hierarchical clustering, Euclidean distance was used as the distance measure and complete-linkage was used to combine clusters. 2-Sample Student t-test was used to find differentially expressed proteins between epilepsy and control groups (per each ROI). We corrected for multiple hypothesis testing by calculating the permutation-based FDR (n=250). Only protein groups with FDR <5% were deemed significantly different between groups (Supplementary Tables 2-8). For each ROI, protein groups absent in a disease group and present in ≥8 cases in the other disease group were considered as differentially expressed. These significant protein groups are included in Supplementary Table 9.

### Pathway and Network Analysis

The genes encoding protein groups significantly altered in epilepsy and control were analyzed using Ingenuity Pathway Analysis (IPA, Qiagen). We analyzed all differentially expressed proteins in the hippocampus (n=777), frontal cortex (249) or dentate gyrus (n=49). We also analyzed differentially expressed proteins in both hippocampus and frontal cortex (n=134) and differentially expressed proteins specific to hippocampus or frontal cortex. All pathway analyses are included in Supplementary Tables 13-18. Network analysis was performed using STRING (v.11). Visualization and analysis of the network was conducted via Cytoscape.

### Cell-type Specificity

For each ROI, genes encoding all significantly altered proteins in epilepsy and controls were assigned cell-type specificity derived from Lake et al. [42], in which each gene had only one cell type annotation from any brain region excluding the cerebellum with 1066 possible annotations. For each ROI, cell type enrichment analysis included the following annotations: neurons, excitatory and inhibitory neurons, astrocytes, endothelial, microglia, and oligodendrocytes. The “neurons” annotation was given to proteins for which the gene in Lake et al. had both “excitatory neuron” and “inhibitory neuron” annotations, with no other cell type associated annotations. For the hippocampus, synaptic proteins were assigned based on the classification of “Synaptic protein” in Lleo et al. [43]. Proteins with a neuron cell type annotation and assigned a synaptic protein were referred to as “neuro-synaptic”.

### Immunohistochemistry (IHC)

To validate proteomics data, we immunostained FFPE tissue sections as described [31]. Sections were incubated overnight at 4°C with either α-SYP antibody (1:50, abcam, catalog #ab8049) or α-TUJ1 antibody (1:300, Biolegend, catalog #802001). Sections were then incubated for 2hr at RT with fluorescent secondary antibodies (All diluted 1:500, Jackson Immunoresearch) and coverslipped (ProLongDiamond Antifade Mountant, Invitrogen). All immunostained sections from one immunostaining combination were scanned at 20X magnification on a NanoZoomer HT2 (Hamamatsu) microscope using the same settings. Fiji was used to analyze images with the same threshold for area positive for pixels relative to total area and reported as a percentage. Statistical analyses used unpaired t-tests. Representative images were obtained by confocal imaging using a Zeiss 700 confocal microscope. For each immunostaining combination, z-stacked images were collected with the same settings at 20X and 63X magnification with a 13 and 11μm z-step and are depicted as a maximum projection image.

## Results

### Quantitative proteomic analysis of epilepsy and non-epilepsy control cases

To identify proteomic alterations in the epilepsy brain, regions of the hippocampus (CA1-3 region) and frontal cortex were microdissected from all epilepsy and control cases (Figure 1A). Cases were matched for sex, age (46 ± 14 years) and formalin archival time (2 ± 1 years) to the best of our ability (Table 1). Histological evaluation of all epilepsy cases identified focal cortical dysplasia (FCD) in 4/14 cases, dysgenesis of the dentate gyrus (DDG) in 4/14 cases and hippocampal sclerosis (HS) in 2/14 cases (Table 1). Control cases had no significant neuropathology (Table 1).

Given the role of the dentate gyrus in epileptogenesis and epileptic neuropathology [44, 45], we microdissected and analyzed the dentate gyrus proteome separately from CA1-3. Given the reduced size of the dentate gyrus, these samples were smaller than hippocampus and frontal cortex samples (4mm^2^ versus 10mm^2^; Figure 1A) and may explain the lower number of protein groups identified and higher coefficient of variation in dentate samples.

Combined, we analyzed 79 samples from 28 cases using label-free quantitative MS and identified 59,903 peptides mapping to 4,894 protein groups (Figure 1B). Additional filtering based on a minimum of 8 cases with quantification per group in a ROI resulted in 3,216 proteins groups being included in our analyses: frontal cortex - 3015, hippocampus - 2990 and dentate gyrus - 2615 proteins. The proteins identified overlapped extensively across regions (Figure 1C).

### Proteomic differences in epilepsy and control cases in hippocampus (CA1-3) and frontal cortex

Protein expression differences between epilepsy and controls were identified in the hippocampus and frontal cortex. Principal Component Analysis (PCA) showed a striking separation of epilepsy and control groups in hippocampus (Student’s t-test, p=2.032×10^−9^, Figure 2A), and to a lesser extent, in frontal cortex (p=0.0125; Figure 2A) in PC1, confirming that the hippocampus is a more impacted region than the frontal cortex. Presence of FCD, DDG and HS did not account for this separation (Figure 2A). Supervised hierarchical clustering revealed robust clustering for all hippocampi and frontal cortices from epilepsy cases, consistent with the PCA analysis (Figure 2B). Variables such as age, sex and neuropathological findings did not drive the clustering (Figure 2B). Only age of epilepsy onset separated different epilepsy cases in the heatmap: cases with an age of onset below or above 20 years old clustered separately. However, several cases did not have comprehensive clinical information. Those cases came from a population with health care disparities that were not under regular medical care and therefore had limited medical records.

**Figure 2.**
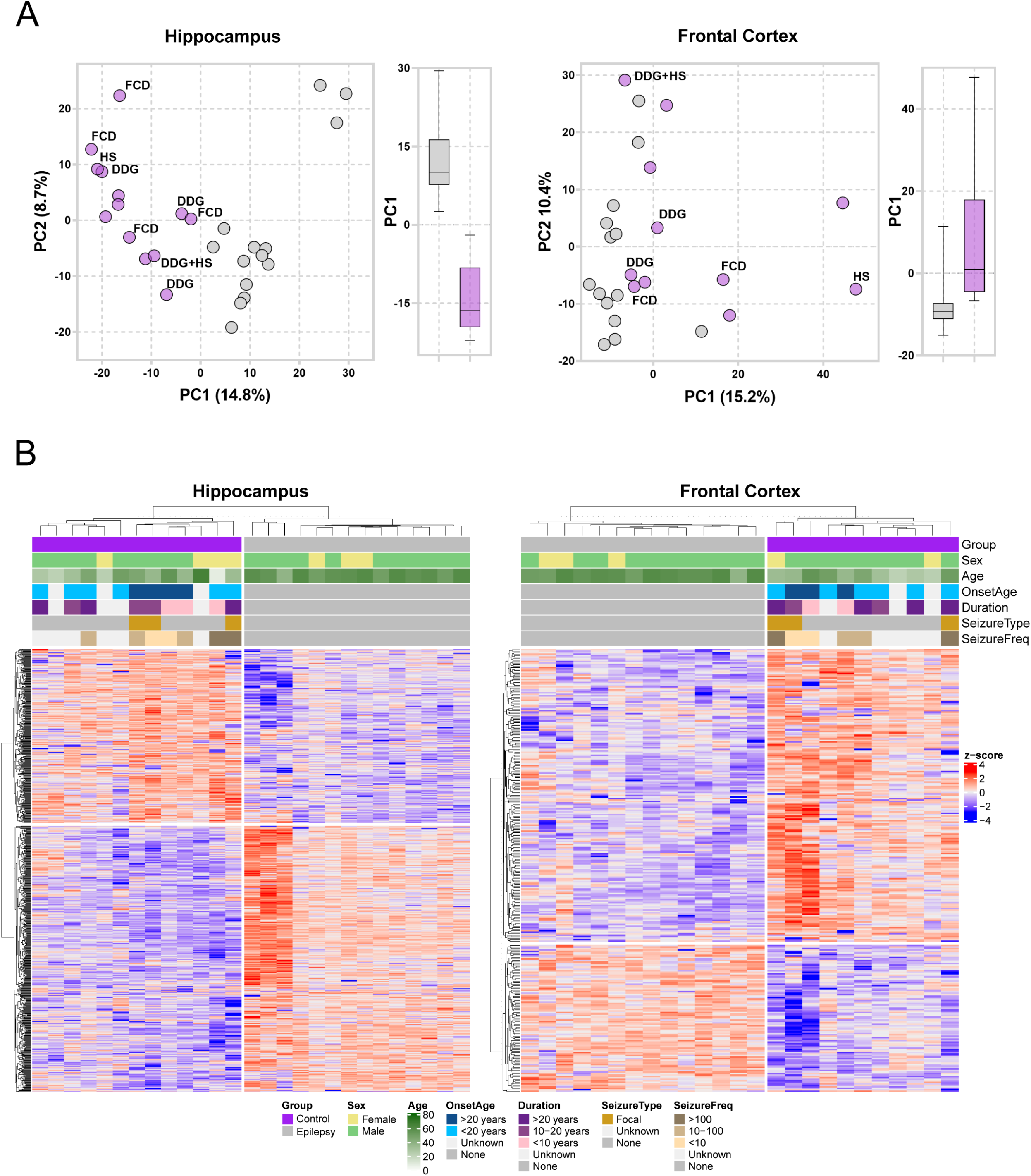
Proteomic differences in epilepsy and control cases in hippocampus (CA1-3) and frontal cortex. **A. PCAs of overall dataset for each region.** Principal component analyses of epilepsy (*purple*) and control (*grey)* cases in the hippocampus (left panel) and frontal cortex (right panel) showing that epilepsy hippocampus (p=2.032×10^−9^) and frontal cortex (p=0.012) contained significantly different protein expression. Each point represents an individual case. Neuropathology (DDG, Dysgenesis of the Dentate Gyrus, FCD; Focal Cortical Dysplasia; HS, Hippocampal Sclerosis) is indicated on the PCA. Significance was determined using a Welch t-test. Error bars represent PC1 min and max. **B. Supervised hierarchical clustering.** Dendrogram showing hierarchical clustering of brain samples based on protein expression levels by region (left panel, hippocampus; right panel, frontal cortex). Information on diagnosis, sex, age, age of disease onset (OnsetAge), disease duration (Duration), seizure type (SeizureType), and frequency of lifetime seizures (SeizureFreq) is indicated according to the legend at the bottom of the graph. Heatmaps on the bottom show scaled (protein z-score) expression values (color-coded according to the legend on the right) for proteins used for clustering.

Differentially expressed proteins between epilepsy and controls were determined using a Student’s t-test and Permutation based 5% FDR. There were significant alterations in the expression of 777 proteins in the hippocampus and 296 proteins in the frontal cortex (Figure 3A-B, Supplementary Tables 4 and 6). 134 proteins were significantly altered in both regions (Supplementary Table 10). The top 20 most significantly altered proteins in epilepsy by region are presented in Table 2 and 3. Of the 40 top altered proteins, 30 (75%) have been associated with epilepsy in humans or animal models (Table 2 and 3). We observed that several altered proteins were encoded by genes in which mutations cause epilepsy [5], including SCN2A, SCN8A, STXBP1, GABBR1, GABBR2, GABRA1, SLC25A12, SLC25A22, SLC6A1, SCARB2 (Figure 3C).

**Figure 3.**
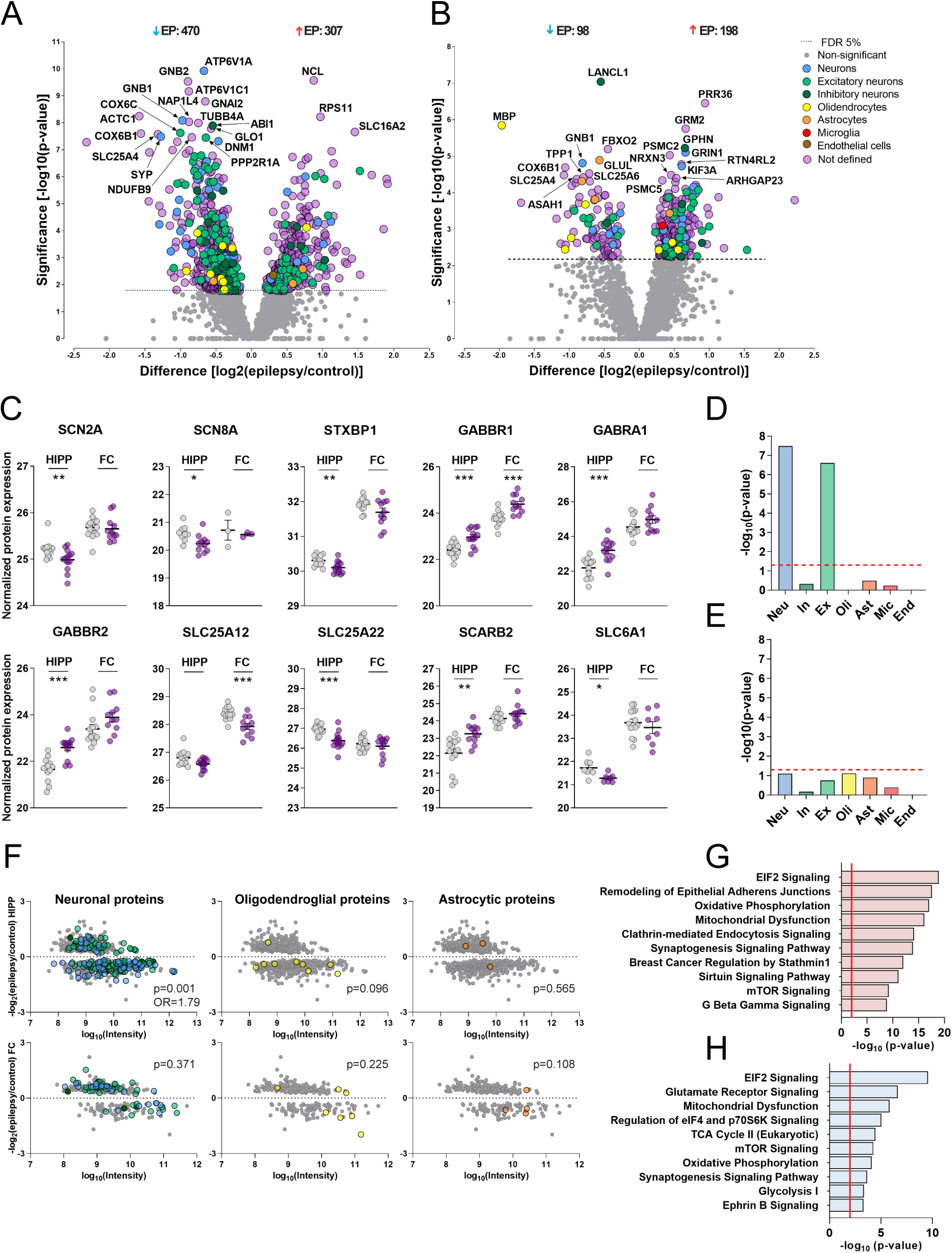
Hippocampal and cortical differences in the epilepsy proteome. **A-B. Protein differential abundance in the hippocampus and frontal cortex.** Volcano plots show protein expression differences of all proteins identified in the hippocampus (A) and frontal cortex (B). All color-coded proteins had significantly altered expression in epilepsy and are indicated above the designated line showing 5% FDR. Proteins were assigned to a specific cell-type (Neu, Neuron [*blue*]; In, Inhibitory Neuron [*dark green*]; Ex, Excitatory Neuron [*light green*]; Oli, Oligodendroglia [*yellow*]; Ast, Astrocyte [*orange*]; Mic, Microglia [*red*]; End, Endothelial cell [*brown*]). Significance was determined using a Student’s t-test with Permutation based False Discovery Rate (FDR) set to 5% (exact p-values for each protein group are provided in Supplementary Table 2). Proteins with higher expression in epilepsy are on the right of the plot and proteins with lower expression are on the left. The 20 top most significantly altered proteins are identified by gene abbreviations. **C. Proteins of interest with significantly altered levels in epilepsy in hippocampus and frontal cortex.** Individual points show protein expression in each individual case. Data shows mean ± SEM; **** p<0.0001; *** p<0.001; ** p<0.01. HIPP = hippocampus, FC = frontal cortex. **D-E. Cell type specific enrichment.** For each brain region, cell type enrichment analysis was performed for each cell type using a Fisher’s exact test. Significance threshold is represented by a dotted red line (p<0.05); **** p<0.0001 **F. Cell type specific protein expression differences in epilepsy.** Each graph plots all significant proteins quantified in the epilepsy hippocampus (top panels) or frontal cortex (bottom panels). The further the point is away from the dashed line at 0, the greater the difference in expression in epilepsy and controls. Proteins with a positive log ratio have greater expression in epilepsy, and proteins with a negative log ratio have a lower expression in epilepsy. All color-coded proteins correspond to neuronal proteins (Neuron, *blue*; Inhibitory Neuron, *dark green*; Excitatory Neuron, *dark green*), oligodendrocyte proteins (*yellow*) and astrocyte proteins (*orange*). The cumulative cell type specific protein differences showed that there were significantly less neuronal proteins in the hippocampus in epilepsy (p=0.001) but not in the frontal cortex (p=0.371). No differences in the amount of oligodendroglial (hippocampus, p=0.096; frontal cortex, p=1), astrocytic (hippocampus, p=0.565; frontal cortex, p=0.108), microglial (not shown) or endothelial proteins (not shown) were observed. Significance was determined using a two-tailed Fisher’s exact test comparing cumulative cell type specific protein expression between epilepsy and control cases. **G-H. Pathway analysis of the altered proteome in the hippocampus and frontal cortex.** IPA was used to identify the top 10 signaling pathways associated with the genes encoding protein groups that were significantly altered in both the hippocampus (top, pink) and the frontal cortex (bottom, blue). The red line corresponds to FDR corrected p-value thresholds (hippocampus, p=0.0001; frontal cortex, p=0.001). All pathways and proteins are shown in Supplementary Tables 13-14.

**Table 2.** Top 20 Significant Proteins in the Hippocampal CA1-3 Region of Epilepsy Cases.

| Gene ID | Protein name | UniProt ID | p Value | Fold Change | Human | Animal model | References |
| --- | --- | --- | --- | --- | --- | --- | --- |
| **<u>Decreased</u>** |  |  |  |  |  |  |  |
| ATP6V1A | V-type proton ATPase catalytic subunit A | P38606 | 1.202E-10 | 1.591 | ☒ | ☒ | Mutations result in developmental delay with epilepsy [84]; protein increased in MTLE hippocampus surgical resections compared to autopsy [85]; protein decreased in childhood cortical dysplasia with epilepsy [86]; gene expression decreased in hippocampus of status epilepticus murine model [87] |
| GNB2 | Guanine nucleotide-binding protein G(I)/G(S)/G(T) subunit beta-2 | P62879 | 2.951E-10 | 1.853 | ☐ | ☒ | Gene expression increased in hippocampus of status epilepticus murine model [87] |
| ATP6V1C1 | V-type proton ATPase subunit C 1 | P21283 | 6.918E-10 | 1.840 | ☐ | ☒ | Gene expression decreased in hippocampus of status epilepticus murine model [87] |
| GNAI2 | Guanine nucleotide-binding protein G(i) subunit alpha-2 | P04899 | 1.622E-09 | 1.569 | ☒ | ☒ | Gene expression increased in MTLE + HS temporal cortex surgical resections compared to autopsy [82]; gene expression decreased in hippocampus of status epilepticus murine model [87] |
| ACTC1 | Actin, alpha cardiac muscle 1 | P68032 | 5.754E-09 | 2.990 | ☐ | ☐ |  |
| GNB1 | Guanine nucleotide-binding protein G(I)/G(S)/G(T) subunit beta-1 | P62873 | 8.128E-09 | 1.959 | ☒ | ☐ | Mutations result in neurodevelopmental delay and seizures [5, 88, 89]; gene expression decreased in MTLE + HS temporal cortex surgical resections compared to autopsy [82] |
| NAP1L4 | Nucleosome assembly protein 1-like 4 | Q99733 | 9.333E-09 | 1.840 | ☒ | ☒ | Gene expression decreased in MTLE + HS temporal cortex surgical resections compared to autopsy [82]; gene expression decreased in hippocampus of status epilepticus murine model [87] |
| TUBB4B | Tubulin beta-4B chain | P68371 | 1.023E-08 | 1.682 | ☐ | ☐ |  |
| ABI1 | Abl interactor 1 | Q8IZP0 | 1.288E-08 | 1.464 | ☐ | ☒ | Gene expression increased in hippocampus of status epilepticus murine model [87] |
| GLO1 | Lactoylglutathione lyase | Q04760 | 1.66E-08 | 1.485 | ☐ | ☒ | Gene expression decreased in hippocampus of status epilepticus murine model [87] |
| COX6C | Cytochrome c oxidase subunit 6C | P09669 | 2.399E-08 | 2.000 | ☐ | ☐ |  |
| COX6B1 | Cytochrome c oxidase subunit VIb isoform 1 | P14854 | 2.512E-08 | 2.949 | ☒ | ☒ | Epilepsy related gene – mitochondrial disease [5]; gene expression decreased in MTLE + HS temporal cortex surgical resections compared to autopsy [82]; gene expression decreased in hippocampus of status epilepticus murine model [87] |
| SLC25A4 | ADP/ATP translocase 1 | P12235 | 2.692E-08 | 2.479 | ☒ | ☒ | Gene expression decreased in MTLE + HS temporal cortex surgical resections compared to autopsy [82]; gene expression decreased in hippocampus of status epilepticus murine model [87] |
| SYP | Synaptophysin | P08247 | 3.236E-08 | 2.428 | ☒ | ☒ | Epilepsy related gene – mental retardation [5, 90]; gene expression decreased in MTLE + HS temporal cortex surgical resections compared to autopsy [82]; gene expression decreased in hippocampus of status epilepticus murine model [87]; protein decreased in hippocampus of status epilepticus rat model [91]; protein decreased in hippocampus of APP/PS1 transgenic mice with seizures [92] |
| NDUFB9 | NADH dehydrogenase [ubiquinone] 1 beta subcomplex subunit 9 | Q9Y6M9 | 3.467E-08 | 1.790 | ☒ | ☒ | Gene expression decreased in MTLE + HS temporal cortex surgical resections compared to autopsy [82]; gene expression increased in hippocampus of status epilepticus murine model [87] |
| PPP2R1A | Serine/threonine-protein phosphatase 2A 65 kDa regulatory subunit A alpha isoform | P30153 | 3.548E-08 | 1.558 | ☒ | ☐ | Epilepsy related gene – mental retardation [5]; protein increased in childhood cortical dysplasia with epilepsy [86] |
| DNM1 | Dynamin-1 | Q05193 | 4.898E-08 | 1.385 | ☒ | ☒ | Mutations result in early infantile epileptic encephalopathy [5, 93, 94]; gene expression decreased in hippocampus of status epilepticus murine model [87] |
| **<u>Increased</u>** |  |  |  |  |  |  |  |
| NCL | Nucleolin | P19338 | 2.692E-10 | 1.828 | ☒ | ☒ | Gene expression decreased in MTLE + HS temporal cortex surgical resections compared to autopsy [82]; protein increased early in hippocampus of status epilepticus rat model [95, 96] |
| RPS11 | 40S ribosomal protein S11 | P62280 | 6.026E-09 | 1.959 | ☐ | ☐ |  |
| SLC16A2 | Monocarboxylate transporter 8 | P36021 | 2.188E-08 | 2.732 | ☒ | ☒ | Gene expression increased in MTLE + HS temporal cortex surgical resections compared to autopsy [82]; gene expression increased in hippocampus of status epilepticus murine model [87] |

MTLE + HS = mesial temporal lobe epilepsy with hippocampal sclerosis

**Table 3.** Top 20 Significant Proteins in the Frontal Cortex of Epilepsy Cases.

| Gene ID | Protein name | UniProt ID | p Value | Fold Change | Human | Animal model | References |
| --- | --- | --- | --- | --- | --- | --- | --- |
| **<u>Decreased</u>** |  |  |  |  |  |  |  |
| LANCL1 | Glutathione S-transferase LANCL1 | O43813 | 9.12E-08 | 1.464 | ☐ | ☒ | Gene expression decreased in hippocampus of status epilepticus murine model [87] |
| MBP | Myelin basic protein | P02686 | 1.445E-06 | 3.891 | ☒ | ☒ | Gene expression decreased in MTLE + HS temporal cortex surgical resections compared to autopsy [82]; gene expression decreased in hippocampus of status epilepticus murine model [87]; protein decreased and myelin damage in hippocampus and cortex of status epilepticus rat models [97, 98] |
| FBXO2 | F-box only protein 2 | Q9UK22 | 6.457E-06 | 1.366 | ☒ | ☒ | Gene expression decreased in MTLE + HS temporal cortex surgical resections compared to autopsy [82]; protein decreased in childhood cortical dysplasia with epilepsy [86]; gene expression decreased in hippocampus of status epilepticus murine model [87] |
| GLUL | Glutamine synthetase | P15104 | 1.288E-05 | 1.485 | ☒ | ☒ | Epilepsy related gene – amino acid metabolic disturbance [5, 99]; protein decreased in childhood cortical dysplasia with epilepsy [86]; protein decreased in MTLE + HS [100]; GLUL knockout mice develop progressive neurodegeneration and seizures [101] |
| GNB1 | Guanine nucleotide-binding protein G(I)/G(S)/G(T) subunit beta-1 | P62873 | 1.549E-05 | 1.753 | ☒ | ☐ | Mutations result in neurodevelopmental delay and seizures [5, 88, 89]; gene expression decreased in MTLE + HS temporal cortex surgical resections compared to autopsy [82] |
| COX6B1 | Cytochrome c oxidase subunit 6B1 | P14854 | 2.042E-05 | 2.085 | ☒ | ☒ | Epilepsy related gene – mitochondrial disease [5]; gene expression decreased in MTLE + HS temporal cortex surgical resections compared to autopsy [82]; gene expression decreased in hippocampus of status epilepticus murine model [87] |
| SLC25A6 | ADP/ATP translocase 3 | P12236 | 2.951E-05 | 1.636 | ☐ | ☐ |  |
| SLC25A4 | ADP/ATP translocase 1 | P12235 | 3.311E-05 | 2.114 | ☒ | ☒ | Gene expression decreased in MTLE + HS temporal cortex surgical resections compared to autopsy [82]; gene expression decreased in hippocampus of status epilepticus murine model [87] |
| TPP1 | Tripeptidyl-peptidase 1 | O14773 | 3.802E-05 | 1.741 | ☒ | ☒ | Epilepsy related gene – neuronal ceroid lipofuscinosis [5]; gene expression increased in MTLE + HS temporal cortex surgical resections compared to autopsy [82]; gene expression increased in hippocampus of status epilepticus murine model [87] |
| ASAH1 | Acid ceramidase | Q13510 | 4.169E-05 | 1.853 | ☒ | ☒ | Epilepsy related gene – developmental disorder of spinal muscular atrophy with progressive myoclonic epilepsy [5, 102]; gene expression increased in MTLE + HS temporal cortex surgical resections compared to autopsy [82]; protein decreased in childhood cortical dysplasia with epilepsy [86]; gene expression increased in hippocampus of status epilepticus murine model [87] |
| **<u>Increased</u>** |  |  |  |  |  |  |  |
| PRR36 | Proline-rich protein 36 | Q9H6K5 | 3.548E-07 | 1.919 | ☐ | ☐ |  |
| GRM2 | Metabotropic glutamate receptor 2 | Q14416 | 1.778E-06 | 1.580 | ☒ | ☒ | Gene expression decreased in MTLE + HS [103] and in models of status epilepticus [104, 105]; GRM2 knockout mice are NMDA toxicity resistant [106] |
| GPHN | Gephyrin | Q9NQX3 | 5.888E-06 | 1.569 | ☒ | ☒ | Epilepsy related gene – metabolic disorder [5]; microdeletion results in seizures and idiopathic generalized epilepsy [107, 108]; abnormally spliced variants in hippocampus of MTLE surgical resections [109]; protein increased in dentate gyrus of chronic status epilepticus rat model [110] |
| GRIN1 | Glutamate receptor ionotropic, NMDA 1 | Q05586 | 7.943E-06 | 1.580 | ☒ | ☒ | Epilepsy related gene – mental retardation [5]; gene expression increased in amygdala of MTLE + HS compared to autopsy [111]; gene expression decreased in hippocampus of status epilepticus murine model [87] |
| PSMC2 | 26S proteasome regulatory subunit 7 | P35998 | 9.333E-06 | 1.347 | ☐ | ☐ |  |
| RTN4RL2 | Reticulon-4 receptor-like 2 | Q7M6Z0 | 1.66E-05 | 1.516 | ☐ | ☐ |  |
| KIF3A | Kinesin-like protein KIF3A | Q9Y496 | 1.862E-05 | 1.526 | ☒ | ☐ | Gene expression decreased in MTLE + HS temporal cortex surgical resections compared to autopsy [82] |
| NRXN3 | Neurexin-3-beta | Q9HDB5 | 3.311E-05 | 1.366 | ☒ | ☐ | Deletions reported in patients with epilepsy and developmental delay [112] |
| ARHGAP23 | Rho GTPase Activating Protein 23 | Q9P227 | 3.981E-05 | 1.444 | ☐ | ☒ | Gene expression decreased in hippocampus of status epilepticus murine model [87] |
| PSMC5 | 26S proteasome regulatory subunit 8 | P62195 | 4.571E-05 | 1.257 | ☒ | ☐ | Gene expression decreased in MTLE + HS temporal cortex surgical resections compared to autopsy [82] |

MTLE + HS = mesial temporal lobe epilepsy with hippocampal sclerosis

We sought to determine whether protein differences in epilepsy were cell type specific. Of the 777 altered proteins in the epilepsy hippocampus, neuronal proteins were significantly enriched (Fisher’s exact test, p=6.49×10^−7^), particularly excitatory neuron proteins (p=7.128×10^−6^) (Figure 3D). There was no enrichment of inhibitory neuron, oligodendroglia, astrocyte, microglia, or endothelial proteins (Figure 3D). These enrichments were not observed in the frontal cortex (Figure 3E), suggesting more extensive protein alterations in hippocampal neurons. All neuron-specific proteins (“neuron”, “inhibitory neuron” or “excitatory neuron” proteins) were predominantly decreased in the epileptic hippocampus compared to controls (Fisher’s exact test, p=0.0013, Figure 3F).

Pathway analysis showed that altered hippocampal and frontal cortex proteins were significantly associated with four pathways: protein biogenesis, mitochondrial function, synaptic plasticity, and G-protein signaling (Figure 3G-H, Supplementary Tables 13-14). Proteins associated with EIF2 signaling were particularly enriched. Most (37/53) were ribosomal proteins and 96.9% were upregulated (19/19 in frontal cortex, 31/33 in the hippocampus), indicating increased translational machinery and ribosome biogenesis. Other proteins involved in this pathway included translation initiation factors (EIF2AK2, EIF3E, EIF3J, EIF4E, EIF4G1) and protein kinases (MAPK1, MAPK3). The mTOR signaling pathway, as well as the regulation of eIF4 and p70S6K signaling pathway - which is part of the mTORC1-S6K signaling axis - were also significantly represented, suggesting that mTOR pathway may drive this increase in proteins related to protein synthesis. The mTOR signaling pathway is involved in protein synthesis, synaptic plasticity, autophagy and other functions which may influence neuronal excitability and epileptogenesis [46, 47]. Oxidative phosphorylation proteins were also enriched in both regions, with broader evidence of altered mitochondrial function including pathways such as mitochondrial dysfunction, sirtuin signaling, glycolysis, and TCA cycle, highlighting impaired neuronal energy regulation in epilepsy. The synaptogenesis signaling pathway was also significantly enriched, confirming synaptic dysfunction resulting from epilepsy. Finally, G-protein signaling was also significantly represented in both regions (G Beta Gamma Signaling/Ephrin B signaling; Figure 3G-H, Supplementary Tables 13-14). Notably, we observed a unique enrichment of glutamate receptor signaling pathway proteins in the frontal cortex and in the remodeling of epithelial adherens junction pathway in the hippocampus. Combined, these results suggest that although changes were more prominent in the hippocampus, similar pathways were enriched in frontal cortex, identifying widespread brain protein alterations in epilepsy.

### Proteomic differences in epilepsy and control cases in the dentate gyrus

Proteomic analysis of the dentate gyrus in epilepsy was limited by smaller tissue area and increased variation. PCA only showed a modest separation between epilepsy and control cases (p=0.081) in the dentate gyrus (Figure 4A). However, quantification of individual proteins showed that 49 proteins were significantly altered in the dentate gyrus in epilepsy (Figure 4B, Supplementary Table 8). Notably, 24 proteins were uniquely altered in the dentate gyrus, including 2 identified only in dentate gyrus (CPSF1, FAM120C) and 2 identified only in dentate gyrus and frontal cortex (CDC5L, AMER2) (Supplementary Table 2). Hierarchical clustering induced by pre-selection of these differently expressed proteins showed that 10/13 epilepsy cases clustered, supporting dentate gyrus dysfunction in epilepsy (Figure 4C). Table 4 lists the 20 most significantly altered proteins; 16 (80%) are associated with epilepsy in humans or animal models, while 4 (20%) were novel (AMER2, ATP5BP, RPL15, TPI1), adding to the list of potential new epilepsy-associated proteins (Table 2 and 3). The G Protein Subunit Beta 1 (GNB1) was the only protein to rank among the top 20 significantly altered proteins in all regions (hippocampus, p= 8.19×10^−9^; frontal cortex; p=1.55×10^−5^, dentate gyrus; p=6.96×10^−6^, Figure 4D). Despite the epileptic encephalopathies that GNB1 mutations can cause [48], the underlying disease mechanisms are still largely unknown and the specific role of GNB1 in epileptogenesis has not yet been studied. Pathway analysis of altered proteins in the dentate gyrus showed the most enrichment in proteins involved in mitochondrial dysfunction, revealing a common epilepsy proteomic signature in all regions (Figure 4E, Supplementary Table 15).

**Figure 4.**
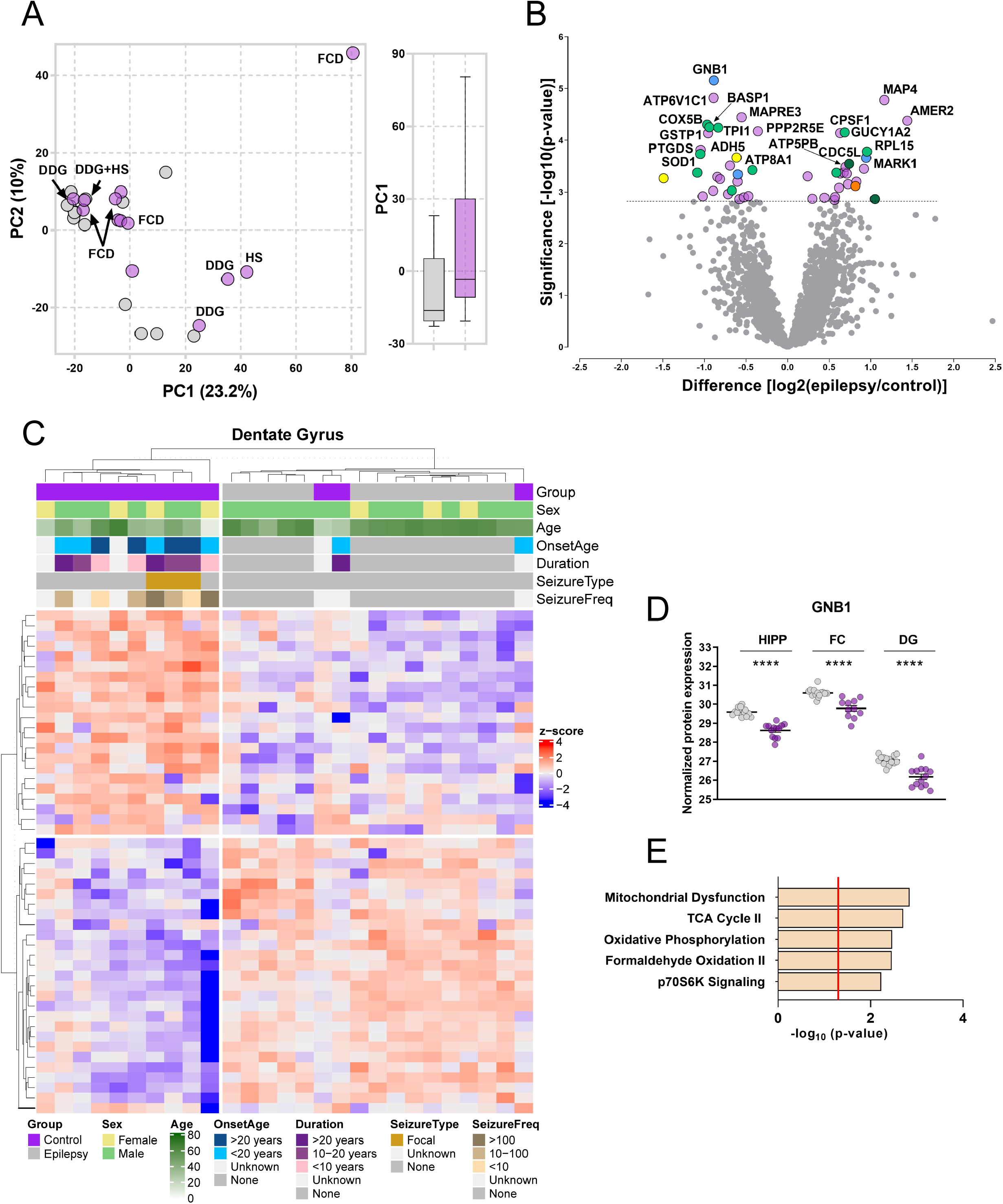
Proteomic differences in epilepsy and control cases in the dentate gyrus. **A. PCA of overall dataset for the dentate gyrus.** The principal component analysis of epilepsy (*purple*) and control (*grey)* cases in the dentate gyrus shows only modest separation between groups (p=0.08). Each point represents an individual case. Neuropathology (DDG, Dysgenesis of the Dentate Gyrus, FCD, Focal Cortical Dysplasia; HS, Hippocampal Sclerosis) is indicated on the PCA. **B. Protein differential abundance in the hippocampus and frontal cortex.** The volcano plot shows protein expression differences of all proteins identified in the dentate gyrus. All color-coded proteins had significantly altered expression in epilepsy. Significance was determined using a Student’s t-test with Permutation based False Discovery Rate (FDR) set to 5% (exact p-values for each protein group are provided in Supplementary Table 2). Proteins with higher expression in epilepsy fall to the right of the plot and proteins with lower expression fall to the left of the plot. The 20 top most significantly altered proteins are identified with gene abbreviations. **C. Supervised hierarchical clustering.** The dendrogram shows hierarchical clustering of brain samples based on protein expression levels in the dentate gyrus. Information on diagnosis, sex, age, age of onset (OnsetAge), disease duration (Duration), Seizure Type (SeizureType), and frequency of lifetime seizure (SeizureFreq) is indicated with color bars according to the legend at the bottom of the graph. The heatmap on the bottom shows scaled (protein z-score) expression values (color-coded according to the legend on the right) for proteins used for clustering. **D. Pathway analysis of the altered proteome in the dentate gyrus.** IPA was performed on genes encoding for protein groups significantly altered in the dentate gyrus. The red line corresponds to FDR corrected p-value threshold (p=0.05). **E. GNB1 widespread downregulation in epilepsy.** Individual points show protein expression in each individual case. Data shows mean ± SEM; **** p<0.0001; HIPP = hippocampus, FC = frontal cortex, DG= dentate gyrus.

**Table 4.** Top 20 Significant Proteins in the Dentate Gyrus of Epilepsy Cases.

| Gene ID | Protein name | UniProt ID | p Value | Fold Change | Human | Animal model | References |
| --- | --- | --- | --- | --- | --- | --- | --- |
| **<u>Decreased</u>** |  |  |  |  |  |  |  |
| GNB1 | Guanine nucleotide-binding protein G(I)/G(S)/G(T) subunit beta-1 | P62873 | 6.918E-06 | 1.853 | ☒ | ☐ | Mutations result in neurodevelopmental delay and seizures [5, 88, 89]; gene expression decreased in MTLE + HS temporal cortex surgical resections compared to autopsy [82] |
| ATP6V1C1 | V-type proton ATPase subunit C 1 | P21283 | 1.514E-05 | 1.866 | ☐ | ☒ | Gene expression decreased in hippocampus of status epilepticus murine model [87] |
| MAPRE3 | Microtubule-associated protein RP/EB family member 3 | Q9UPY8 | 3.631E-05 | 1.464 | ☐ | ☒ | Gene expression decreased in hippocampus of status epilepticus murine model [87] |
| COX5B | Cytochrome c oxidase subunit 5B, mitochondrial | P10606 | 5.012E-05 | 1.959 | ☒ | ☐ | Gene expression decreased in MTLE + HS temporal cortex surgical resections compared to autopsy [82] |
| BASP1 | Brain acid soluble protein 1 | P80723 | 5.623E-05 | 1.919 | ☒ | ☒ | Gene expression decreased in MTLE + HS temporal cortex surgical resections compared to autopsy [82]; gene expression decreased in hippocampus of status epilepticus murine model [87]; protein decreased in hippocampus of status epilepticus murine model [26] |
| TPI1 | Triosephosphate isomerase | P60174 | 5.754E-05 | 1.778 | ☐ | ☐ |  |
| PPP2R5E | Serine/threonine-protein phosphatase 2A 56 kDa regulatory subunit epsilon isoform | Q16537 | 6.761E-05 | 1.283 | ☒ | ☐ | Gene expression decreased in MTLE + HS temporal cortex surgical resections compared to autopsy [82] |
| GSTP1 | Glutathione S-transferase P | P09211 | 7.413E-05 | 1.945 | ☒ | ☐ | Gene expression increased in MTLE + HS temporal cortex surgical resections compared to autopsy [82]; protein increased in MTLE hippocampus surgical resections compared to autopsy [85] |
| PTGDS | Prostaglandin-H2 D-isomerase | P41222 | 0.0001549 | 2.056 | ☒ | ☒ | Gene expression increased in MTLE + HS temporal cortex surgical resections compared to autopsy [82]; protein decreased in childhood cortical dysplasia with epilepsy [86]; gene expression decreased in hippocampus of status epilepticus murine model [87]; gene expression increased in hippocampus of status epilepticus rat model [113] |
| SOD1 | Superoxide dismutase [Cu-Zn] | P00441 | 0.000182 | 2.071 | ☒ | ☒ | Gene expression decreased in MTLE + HS temporal cortex surgical resections compared to autopsy [82]; protein decreased in CSF of patients with TLE [114]; gene expression decreased in hippocampus of status epilepticus murine model [87] |
| ATP8A1 | Phospholipid-transporting ATPase IA | Q9Y2Q0 | 0.0002138 | 1.526 | ☐ | ☒ | Gene expression increased in hippocampus of status epilepticus murine model [87]; gene expression decreased in hippocampus of status epilepticus transgenic murine model [115] |
| ADH5 | Alcohol dehydrogenase class-3 | P11766 | 0.000309 | 1.613 | ☒ | ☐ | Protein decreased in childhood cortical dysplasia with epilepsy [86] |
| <u>**Increased**</u> |  |  |  |  |  |  |  |
| MAP4 | Microtubule-associated protein 4 | P27816 | 1.66E-05 | 2.235 | ☒ | ☒ | Protein increased in childhood cortical dysplasia with epilepsy [86]; gene expression increased in hippocampus of status epilepticus murine model [87] |
| AMER2 | APC membrane recruitment protein 2 | Q8N7J2 | 4.169E-05 | 2.713 | ☐ | ☐ |  |
| GUCY1A2 | Guanylate cyclase soluble subunit alpha-2 | P33402 | 7.079E-05 | 1.613 | ☒ | ☐ | Gene expression decreased in MTLE + HS temporal cortex surgical resections compared to autopsy [82] ; gene expression increased in hippocampus of status epilepticus murine model [87] |
| CPSF1 | Cleavage and polyadenylation specificity factor subunit 1 | Q10570 | 7.244E-05 | 1.548 | ☐ | ☐ |  |
| RPL15 | 60S ribosomal protein L15 | P61313 | 0.000166 | 1.945 | ☐ | ☒ | Protein increased early in hippocampus of status epilepticus rat model [95, 96] |
| MARK1 | Serine/threonine-protein kinase MARK1 | Q9P0L2 | 0.0002188 | 1.932 | ☒ | ☒ | Gene expression decreased in MTLE + HS temporal cortex surgical resections compared to autopsy [82]; gene expression decreased in hippocampus of status epilepticus murine model [87] |
| CDC5L | Cell division cycle 5-like protein | Q99459 | 0.0002884 | 1.670 | ☐ | ☒ | Gene expression decreased in hippocampus of status epilepticus murine model [87] |
| ATP5PB | ATP synthase F | P24539 | 0.0003311 | 1.625 | ☐ | ☐ |  |

MTLE + HS = mesial temporal lobe epilepsy with hippocampal sclerosis, CSF = cerebrospinal fluid

### Inter-regional epilepsy proteomic signature

The common enriched pathways in the hippocampus and frontal cortex (Figure 3G-H), as well as the overlapping altered proteins (Figure 5A), indicate shared protein changes across both regions. We performed correlation analyses of the directional changes in hippocampus and frontal cortex for the (1) 939 significantly altered proteins and (2) 134 common proteins altered in both regions (Figure 5B-C, Supplementary Table 10). We found a modest positive correlation between protein levels in the hippocampus and frontal cortex when all significant proteins were examined (Pearson’s correlation, r^2^=0.3156, Figure 5B), suggesting that the hippocampus was more severely and/or differently impacted compared to the frontal cortex. The correlation improved when analyzing the changes of the 134 significantly different proteins shared by both the hippocampus and frontal cortex (r^2^=0.7302, Figure 5C), confirming broad similar protein alterations throughout the epileptic brain. Interestingly, this correlation was also present in the dentate gyrus (hippocampus/dentate gyrus, r^2^=0.6431; frontal cortex/dentate gyrus, r^2^=0.7163, Figure 5D-E), despite only 13/134 of these proteins being significantly altered in the dentate gyrus (Figure 5A, D-E, Supplementary Table 10), supporting an epilepsy-associated proteomic signature. Hierarchical clustering of these specific proteins show almost complete clustering of all epilepsy cases throughout the three regions analyzed (Figure 5F). These 134 enriched proteins relate to EIF2 and mTOR signaling pathways, mitochondrial dysfunction and other related metabolic pathways such as oxidative phosphorylation, glycolysis, TCA cycle and gluconeogenesis (Figure 5G-H, Supplementary Table 16). Proteins involved in G-protein signaling were also significantly enriched (Ephrin B signaling; Figure 5G-H, Supplementary Table 16).

**Figure 5.**
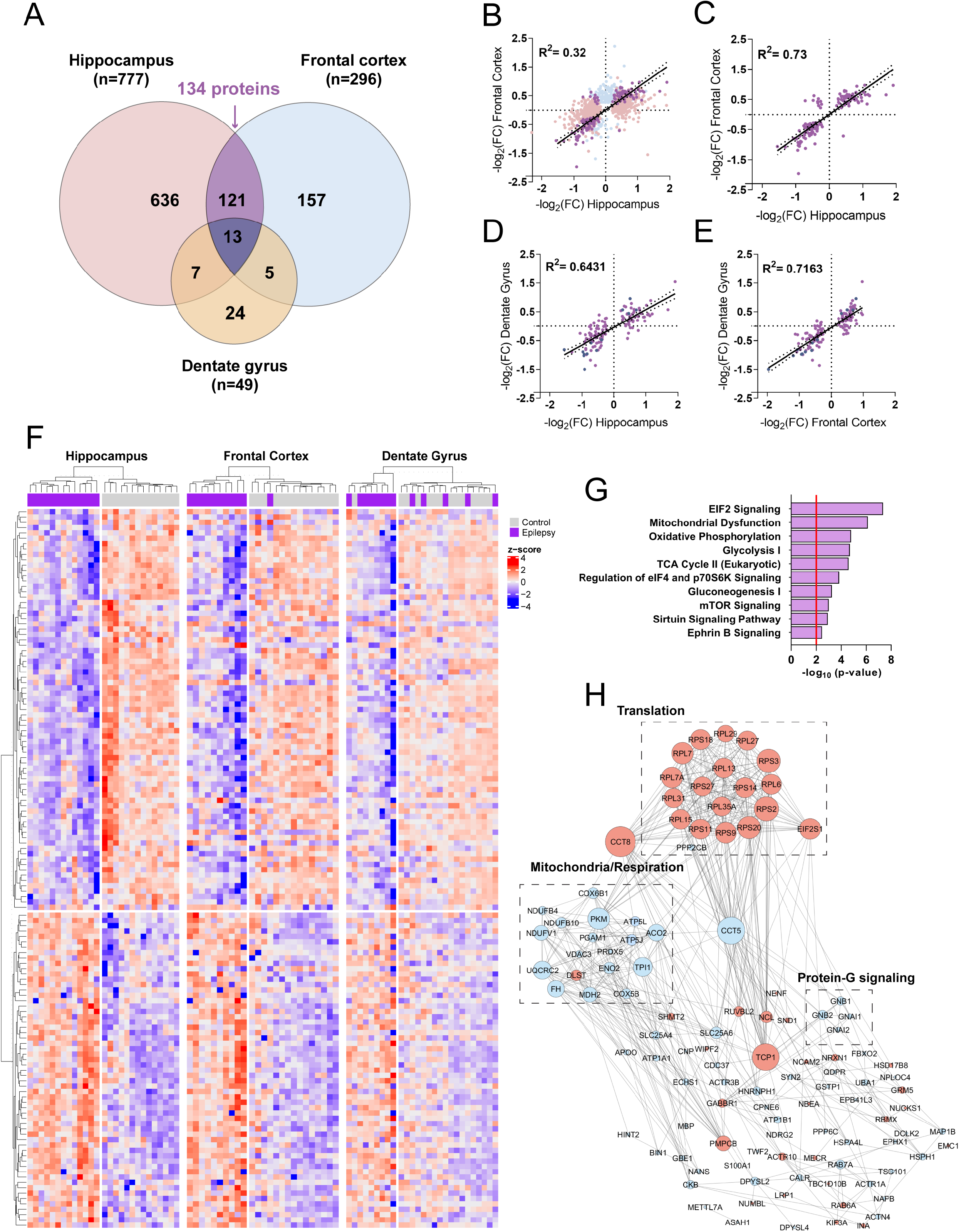
Inter-regional epilepsy proteomic signature. **A. Venn diagram of significant proteins in each region.** The Venn diagram shows the 963 altered proteins in epilepsy across all regions analyzed. 939 proteins were altered in hippocampus and frontal cortex altogether. There was 134 overlapping proteins between the hippocampus and the frontal cortex (*purple*), and 13 overlapping proteins across the three regions (*dark blue*). **B-E. Correlation between the protein fold changes in the hippocampus and the frontal cortex.** Graphs represent correlations using the 939 significantly altered proteins (top left panel, Pearson’s correlation, r^2^=0.32)), and the overlapping 134 proteins altered in both hippocampus and frontal cortex (top right panel, Person’s correlation (r^2^=0.73)). The proteomic signature was also preserved in the dentate gyrus, despite only 13/134 proteins being significant (bottom panels, Pearson’s correlation; hippocampus/dentate gyrus (bottom left panel), r^2^=0.6431; frontal cortex/dentate gyrus (bottom right panel), r^2^=0.7163). **F. Hierarchical clustering of the 134 proteins showing similar directional changes in hippocampus CA1-3, frontal cortex and dentate gyrus.** The dendrogram shows hierarchical clustering of brain samples based on protein expression levels by region. The heatmap on the bottom show scaled (protein z-score) expression values (color-coded according to the legend on the right) for proteins used for clustering. **G. Pathway analysis of the epilepsy proteomic signature.** IPA was performed on genes encoding for protein groups significantly altered in the proteomic signature. The red line corresponds to FDR corrected p-value threshold (p=0.05). **H. Epilepsy proteomic signature protein network.** The network depicts altered proteins in both hippocampus and frontal cortex. Visualization and analysis of the network was conducted via Cytoscape. Color-coding represents directional changes (upregulated shown in *red*, downregulated shown in *blue*). Node sizes correspond to degree of interaction. Disconnected nodes in the network were removed. Black rectangles correspond to proteins involved in pathways of interest.

Together, these results support a broad impact of epilepsy on mitochondrial function, protein biogenesis and G-protein signaling. Notably, synaptic plasticity related signaling pathways were not among shared significant proteins changes across hippocampus and frontal cortex, suggesting that the synaptic-associated mechanisms differ between these regions.

### Intra-regional epilepsy proteomic changes

We found that 635 proteins reached significance in the hippocampus and not in the frontal cortex. These highly significant hippocampal differences were enriched in proteins involved in the remodeling of epithelial junction (p=7.71×10^−18^), with prominent involvement of cytoskeleton-associated proteins (actin-related [ACTC1, ACTG1, ACTN1, ACTR2, ACTR3, ARPC4, ARPC1A], dynamins [DNM1, DNM2. DNM3, DNM1L] and tubulins [TUBA1A, TUBA1B, TUBA4A, TUBB3, TUBB, TUBB2A, TUBB4A]) and clathrin-mediated endocytosis signaling (p=7.92×10^−16^), suggesting a reorganization of the cortical microarchitecture and a remodeling of neuronal circuitries, or an impact on the brain vasculature (Supplementary Table 17). A significant alteration in synaptogenesis signaling pathway proteins was also observed (p=6.26×10^−14^, Supplementary Table 17). These findings, together with the significant decrease of neuronal proteins observed previously (Figure 3F) suggest neuronal synaptic dysfunction in the hippocampus. Neuronal loss is another possible explanation but was not seen histologically. Using a publicly available dataset of the most updated synaptic proteome [43], we confirmed that the significant proteins in the hippocampus were highly enriched in neuro-synaptic proteins (33/63 neuro-synaptic proteins, Fisher’s exact test, p=3.939×10^−6^; Supplementary Table 4). The expression of significant synaptic proteins in epilepsy was dramatically decreased (55/58 proteins, Fisher’s exact test, 8.133×10^−10^, Figure 6A), suggesting extensive synaptic alteration in the hippocampus. These proteomic changes were not significant in the frontal cortex ((Fisher’s exact test, p=0.089; Figure 6B, Supplementary Table 4). We validated the decrease of the key synaptic protein synaptophysin (SYP) in the epileptic hippocampus using IHC (Figure 6C, E, and G). To determine whether these protein changes were unique to this region or simply less pronounced in the frontal cortex than in the hippocampus, we examined the directional changes of the significantly altered hippocampal proteins and their respective directional change in the frontal cortex. When we closely examined these differences, we discovered that the vast majority of these proteins were showing trends for similar changes in the frontal cortex (462/635 proteins in agreement [72%], Fisher’s exact test, p<2.2×10^−16^, Figure 6I), suggesting that overall the same pathological protein changes were happening in the frontal cortex, but not to the same extent. IPA confirmed that the proteins that were in agreement were involved in the remodeling of epithelial adherens junctions (p=2.14×10^−12^), and synaptogenesis (p=1.52×10^−10^). The significantly decreased SYP immunoreactivity in the frontal cortex also supported this observation (Figure 6D, F and H).

**Figure 6.**
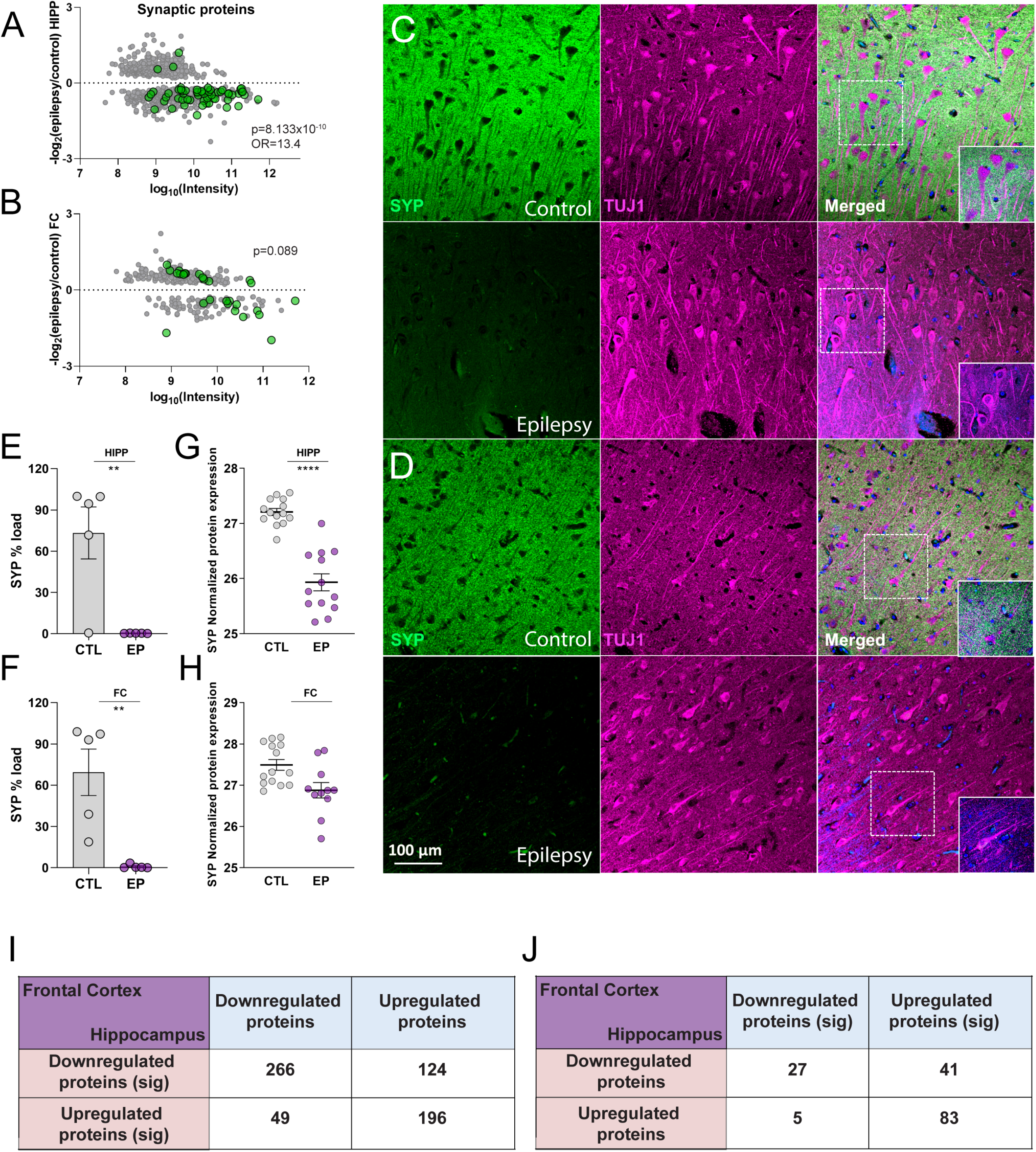
Altered synaptic transmission in epilepsy. **A. The graph depicts all significant proteins identified in the hippocampus.** The further the point is away from the dashed line at 0, the greater the difference in expression in epilepsy and controls. Proteins with a positive log ratio have greater expression in epilepsy and proteins with a negative log ratio have a greater expression in controls. All proteins highlighted in *green* correspond to synaptic proteins. **B.** The graph shows all significant proteins identified in the frontal cortex. All proteins highlighted in *green* correspond to synaptic proteins. **C.** Synaptophysin (SYP) and Tubulin-β 3 (TUJ1) double fluorescent immunohistochemistry of n=5 epilepsy and n=5 control cases in the hippocampus showed major decrease in SYP immunoreactivity in the epileptic hippocampus (panels show CA1 region, images taken at 20X (top panels) and 63X magnification (bottom panels)). **D.** Synaptophysin (SYP) and Tubulin-β 3 (TUJ1) double fluorescent immunohistochemistry of n=5 epilepsy and n=5 control cases in the frontal cortex also showing major decrease in SYP immunoreactivity in the epileptic cortex (images taken at 20X (top panels) and 63X magnification (bottom panels)). **E-H.** Quantification of SYP immunohistochemistry (E-F) and proteomic expression levels (G-H) in the hippocampus CA1-3 (E-G) and frontal cortex (F-H) (n=5/group for IHC and n=14/group for proteomics) epilepsy. For IHC quantification, individual points show SYP % load in each individual case (E-F). Significance was determined using a two-tailed unpaired t-test. Data shows mean ± SD; ** p<0.01. For proteomics, individual points show protein expression in each individual case (G-H). Data shows mean ± SEM; **** p<0.0001. HIPP = hippocampus, FC = frontal cortex. **I-J. Topographical changes in epilepsy support the idea of selective vulnerability in hippocampus.** The left panel shows a contingency table of the protein fold changes in the hippocampus and frontal cortex using the significant proteins in hippocampus only. The right panel shows a contingency table of the protein fold changes in the hippocampus and frontal cortex using the significant proteins in frontal cortex only. Proteins were annotated as either increased or decreased in epilepsy. Only proteins present in both regions were included in the analysis. Significance was determined using a Fisher’s exact test.

Similarly, we found that 156 proteins reached significance in the frontal cortex and not in the hippocampus. While these frontal cortex protein changes were also suggestive of altered synaptogenesis, these changes were mostly enriched in proteins involved in increased glutamate receptor signaling (8/57 proteins, p=2.61×10^−7^, z-score=2.236, Supplementary Table 18), which occurs in glutamatergic synapses, and plays a critical role in excitotoxic damage in epilepsy [49]. Proteins in this pathway included protein Glutamine Synthetase (GLUL), Excitatory amino acid transporters 1 and 2 (SLC1A2, SLC1A3), Glutamate receptor ionotropic NMDA 1 and 2B (GRIN1, GRIN2B), Glutamate receptor ionotropic AMPA 2 (GRIA2) as and Glutamate metabotropic receptor 2 (GRM2). Interestingly, GLUL downregulation was associated with reduced astrocyte protection against glutamate excitotoxicity to neurons [50]. Here again, we found that the majority of these significant changes which seemed specific to the frontal cortex were showing trends for similar directional changes in the hippocampus (110/156 proteins in agreement [72%], p=1.753×10^−7^, Figure 6J), suggesting that most of these changes were also occurring in the hippocampus, yet to a lesser extent. IPA confirmed that the proteins in agreement were most enriched in the glutamatergic signaling pathway (p=7.58×10^−5^).

Combined, our results indicate that 696/939 (74%) significant proteins had similar directional changes in both the hippocampus and frontal cortex in epilepsy (Supplementary Table 11). These findings suggest that most pathological protein changes in epilepsy occurred in both regions but to a different degree, supporting greater vulnerability or a primary role of the hippocampus compared to the frontal cortex.

## Discussion

In this study we have generated the first comprehensive characterization of the human epileptic proteome. We identified extensive protein alterations in epilepsy in multiple brain regions. The epileptic hippocampus and frontal cortex showed dysregulation in pathways associated with mitochondrial function, protein synthesis, synaptic transmission and remodeling of the neuronal network architecture. These protein alterations were present despite the lack of major neurodegeneration. Our proteomics analysis provides a network-level analysis of protein changes in epilepsy and identifies hundreds of new potential drug targets. Given the paucity of comprehensive proteomics studies, it was difficult to cross-validate our findings. However, Wang and colleagues presented the most updated summary of the genes associated with epilepsy, grouping these genes according to the manifestation of epilepsy in specific phenotypes. We compared our list of 939 altered proteins (hippocampus and frontal cortex) to this list and found that 20/939 proteins were encoded by genes in which mutations cause epilepsy [5]. Importantly, 98/939 proteins were encoded by neurodevelopment-associated epilepsy genes, epilepsy-related genes, or potentially epilepsy-associated genes. Our results also suggest that the G-protein heterotrimeric complexes may have a particularly important role in epilepsy and specifically highlighted the widespread decrease in GNB1 expression levels in epilepsy.

Our results showed that the hippocampus was particularly vulnerable in epilepsy and suggested that protein changes in epilepsy may progressively extend through the brain in a topographic manner. The increased vulnerability of the hippocampus was evident in the broad range of epilepsy syndromes included in this study. The hippocampus is an epileptogenic brain region that is highly susceptible to damage from seizures. As a result, hippocampal sclerosis is a common neuropathological feature of epilepsy and its incidence varies from 30-45% of all epilepsy syndromes and 56% of MTLE neuropathologically [51]. Hippocampal sclerosis is the most common pathology associated with drug-resistant epilepsy in older adults [52]. Our cohort only included 2 cases with hippocampal sclerosis, which alone was not enough to explain such extensive hippocampal neuronal alterations. In contrast to earlier studies, we included a heterogeneous case cohort to identify protein alterations across a spectrum of epilepsy syndromes. Our study suggests that hippocampal vulnerability is a consistent feature across all types of epilepsy. It remains unknown why the hippocampus is so vulnerable in epilepsy as well as in ischemia, limbic encephalitis, multiple sclerosis, and neurodegenerative diseases such as Alzheimer’s disease (AD) [53–57]. The cognitive deficits in temporal lobe epilepsy are associated with Alzheimer-like amyloid β and tau protein changes [58]. Hippocampus vulnerability in epilepsy and other neurological disorders may result from a low threshold for glutamatergic excitotoxicity or for synaptic remodeling. Some studies show vulnerability correlates with increased neuroplasticity, which could reflect different energy production mechanisms and requirements, making it highly sensitive to metabolic and oxidative stress, which can occur in epilepsy [59].

Our results suggest that epilepsy affects brain areas in parallel yet to a different degree, supporting that epilepsy may be a progressive disease [60]. Hippocampal protein alterations reflected pathologic changes of degeneration, such as neuronal and synaptic remodeling, which were also observed in the frontal cortex without reaching a significant threshold of 5% FDR (Figure 7). While decreased synaptophysin immunoreactivity was found in temporal lobe epilepsy-associated HS [61], our study is the first to report significantly decreased synaptophysin levels in epilepsy cases in the frontal cortex and hippocampus without hippocampal sclerosis. In contrast, frontal cortex protein changes seemed to mainly reflect earlier pathological changes such as an altered glutamatergic system, with non-significant but similar directional changes in the hippocampus. However, both regions showed mitochondrial dysfunction and altered protein synthesis, although these changes were more prominent in the hippocampus (Figure 7). The shared regional changes in mitochondrial function and protein biosynthesis, as well as the more localized frontal changes in the glutamatergic system suggest that these may be relatively earlier events in the disease process. The lack of significant glutamatergic changes in the hippocampus may reflect the statistical noise of the large protein changes, or that as the disease progresses, these neurons may be selectively lost or compensatory changes wash out any effect. Glutamatergic hyperactivity may be an early insult, driving or resulting from epileptic activity, and leading to oxidative stress and mitochondrial dysfunction; although, this remains unproven. Mitochondrial dysfunction lowers intracellular levels of ATP and alters calcium homeostasis, and these two pathways may be linked in a feed-forward pathologic cycle. [62] Whether mitochondrial dysfunction is a cause or a consequence of epilepsy – or both - our study shows it is a widespread change in the epilepsy brain. Many metabolic and mitochondrial disorders cause seizures and disorders such as myoclonic epilepsy with ragged red fibers [63, 64]. Seizures occur in 35-60% of individuals with primary mitochondrial diseases [65]. Mitochondrial oxidative stress occurs in temporal lobe epilepsy [66–68], but it remains uncertain if this is part of epileptogenesis or a secondary effect of seizures, ASDs or other factors. Mitochondrial functions can impact on neuronal hyperexcitability via generation of ATP, metabolite/neurotransmitter biosynthesis, calcium homeostasis, reactive oxygen species production and apoptosis. Our study suggests that mitochondrial dysfunction may both contribute to and result from epilepsy. Our study also identified significantly increased protein biosynthesis in epilepsy. While this increase could be a result of loss of neuronal homeostasis due to mitochondrial dysfunction [69], increased protein synthesis may be driven by altered mTOR signaling, which was a prominent finding in our results. mTOR signaling pathways are implicated in a number of epileptogenic processes [70]. Our results suggest increased ribosome biogenesis and protein production in chronic epilepsy. Protein synthesis may shift towards greater production of both pro-epileptic factors and compensatory anti-epileptic factors [71]. Also, dysregulated mTOR signaling, part of the proteomic signature of epilepsy, may stimulate excessive synthesis of ion channels and receptors, leading to hyperexcitability [72].

**Figure 7.**
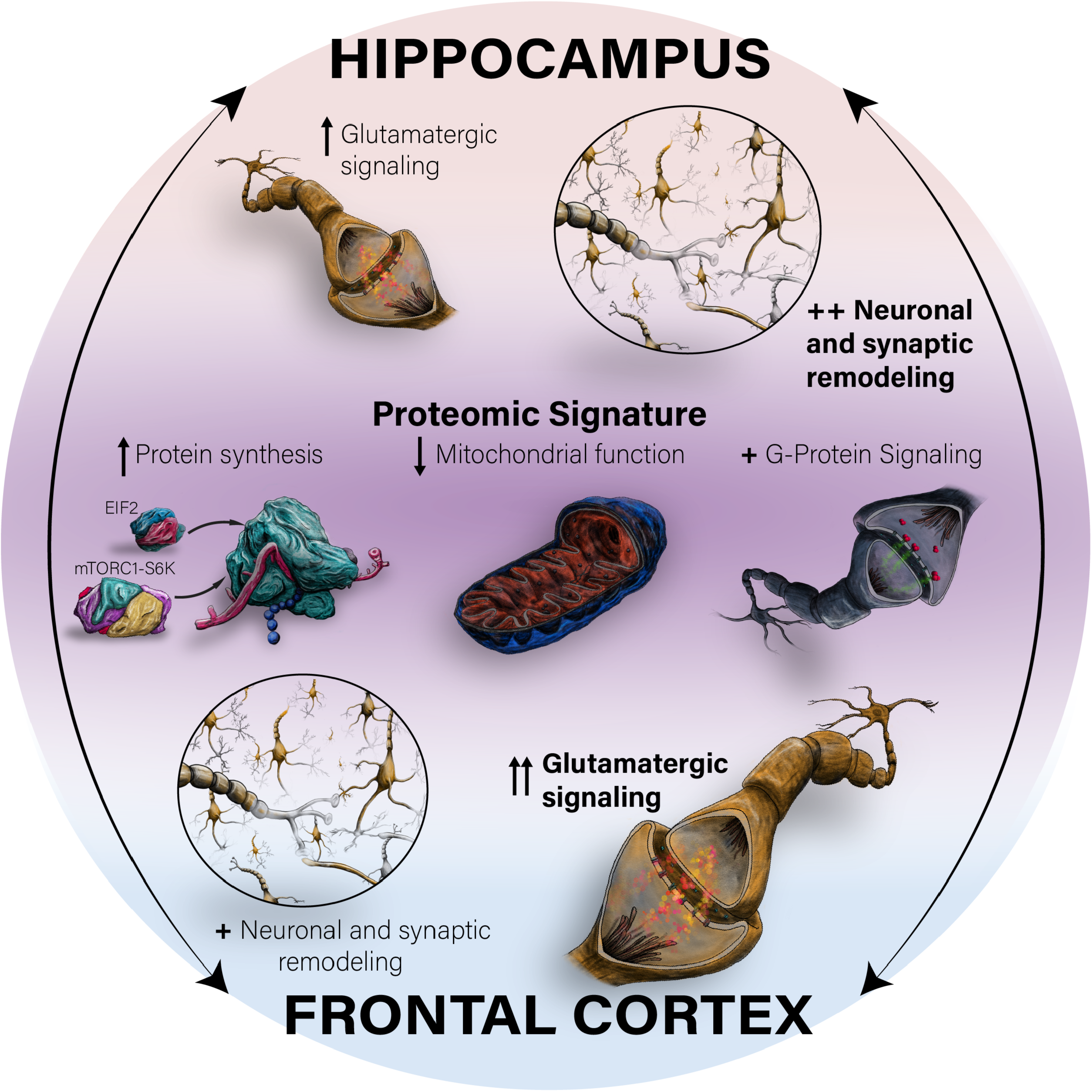
Proteomic alterations in the hippocampus and frontal cortex of epilepsy. The schematic shows broad alterations in both the hippocampus and frontal cortex with the following pathways being significantly altered: protein synthesis through EIF2 and mTORC1-S6K signaling, mitochondrial function, and G-protein signaling (noted as the “Proteomic Signature”). Further alterations include neuronal and synaptic remodeling, which was significant only in the hippocampus however it was also observed in the frontal cortex to a lesser extent. Glutamatergic signaling was specifically significant in the frontal cortex, and it was showing similar but not significant changes in the hippocampus. ↑ increased; ↓ decreased; ++ significantly altered; + altered. Number of symbols represents the magnitude of the alteration.

Our findings identified significant G-protein subunit alterations in epilepsy (Figure 7). Specifically, we detected 20 protein G-subunits in our study, with 13 of them being differentially expressed in at least one region (GNG7, GNAO1, GNG2, GNG4, GNB1, GNB2, GNAI1, GNAI2, GNAQ, GNAZ, GNB4, GNA11, and GNL1). Heterotrimeric G proteins consist of two functional units, an α-subunit (Gα) and a tightly associated βγ complex. After activation, GPCRs change their conformation and transduce this into intracellular signals involved in diverse signaling pathways including the cAMP/PKA pathway, calcium/protein kinase C (PKC) pathway, IP3/DAG/phospholipase C pathway, β-arrestin pathway, protein tyrosine kinase pathway, ERK/MAPK pathway, PI3K/AKT pathway, Rho pathway and G-protein-gated calcium channels inwardly rectifying potassium channels [73]. Animal studies suggests that GPCRs are important mediators of neuronal excitability, supporting a potential therapeutic role in epilepsy [74]. However, no FDA-approved ASD suppresses epileptic seizures by directly acting on GPCRs. GNB1 may be central to GPCR dysfunction in epilepsy [74]. We found that one G-protein subunit, GNB1, was significantly decreased in all the regions. Missense, splice-site and frameshift variants in the GNB1 gene are associated with multiple neurological phenotypes, including seizures in most cases [75–80]. Mice harboring the GNB1 K78R human pathogenic recapitulate many clinical features, including developmental delay, motor and cognitive deficits, and absence-like generalized seizures, and that cellular models displayed extended bursts of firing followed by extensive recovery periods [81]. Given the therapeutic potential of targeting G-proteins, and the novelty of our finding, it would be interesting to further investigate the role of GNB1 in epilepsy. The involvement of GNB1 variants in human epilepsies and our data suggests this gene may play an important role in many epilepsies.

Our study provides a rich resource for epilepsy research, adding to a recent transcriptomic analysis of human MTLE brain [82]. Combining different “-omics” may provide complimentary facets to a more comprehensive understanding of epilepsy pathogenesis and help identify novel therapeutic targets. A limitation of this study is the heterogeneity of our cohort with diverse epilepsies, therapies and genetic backgrounds. However, our study was designed to identify common protein changes in epilepsy across heterogeneous cases which increased variation, making the robust protein changes we observed more significant. Future studies should examine specific epilepsy syndromes as well as the role of ASDs. Our study was not powered to assess the impact of treatments, which could bias this study by altering protein levels and altering non-epilepsy pathways. For example, altered GABAergic receptors or mitochondrial function could reflect ASDs such as clobazam, clonazepam or valproic acid [83].

In conclusion, our study is the first comprehensive proteomic study in the human epilepsy brain and identifies critical pathways novel proteins that may be considered as potential drug targets. These pathways mainly include protein biogenesis, mitochondrial function, and synaptogenesis signaling. Combined, our findings provide a valuable resource for epilepsy research.

## Acknowledgements

This work was supported by FACES and the NIH grant U01 NS090415. This work was supported by funding from the Bluesand Foundation to ED, Philippe Chatrier Foundation to GP. The proteomics work was in part supported by the NYU School of Medicine and a shared instrumentation grant from the NIH, 1S10OD010582-01A1 for the purchase of an Orbitrap Fusion Lumos.

## Supplementary Table Legends

**Supplementary Table 1:** Description of supplementary table columns in Supplementary Tables 2-12.

**Supplementary Table 2:** Overall protein differences in all brain regions of epilepsy and control cases.

**Supplementary Table 3:** All proteins detected in the hippocampal CA1-3 region of epilepsy and control cases.

**Supplementary Table 4:** Significant proteins (777) detected in the hippocampal CA1-3 region of epilepsy and control cases.

**Supplementary Table 5:** All proteins detected in the frontal cortex of epilepsy and control cases.

**Supplementary Table 6:** Significant proteins (296) detected in the frontal cortex of epilepsy and control cases.

**Supplementary Table 7:** All proteins detected in the dentate gyrus of epilepsy and control cases.

**Supplementary Table 8:** Significant proteins (49) detected in the dentate gyrus of epilepsy and control cases.

**Supplementary Table 9:** Proteins detected only in either epilepsy or control cases (12)

**Supplementary Table 10:** Proteins significant (134) in both the hippocampal CA1-3 region and frontal cortex.

**Supplementary Table 11:** Proteins altered in the same direction (up/down) in the hippocampal CA1-3 region and frontal cortex.

**Supplementary Table 12:** Protein differences in all brain regions of epilepsy and control cases, with the addition of imputed data.

**Supplementary Table 13:** Pathway Analysis data from IPA of significant proteins in the hippocampus in epilepsy

**Supplementary Table 14:** Pathway Analysis data from IPA of significant proteins in the frontal cortex in epilepsy

**Supplementary Table 15:** Pathway Analysis data from IPA of significant proteins in the dentate gyrus in epilepsy

**Supplementary Table 16:** Pathway Analysis data from IPA of significant proteins included in the epilepsy proteomic signature

**Supplementary Table 17:** Pathway Analysis data from IPA of significant proteins only in the hippocampus in epilepsy

**Supplementary Table 18:** Pathway Analysis data from IPA of significant proteins only in the frontal cortex in epilepsy

